# A uniform stress response of stream microbiomes in the hyporheic zone across North America

**DOI:** 10.1101/2025.02.16.638492

**Authors:** Tom L. Stach, Jörn Starke, Feriel Bouderka, Till L. V. Bornemann, André R. Soares, Michael J. Wilkins, Amy E. Goldman, James C. Stegen, Mikayla A. Borton, Alexander J. Probst

**Affiliations:** Environmental Metagenomics, Research Center One Health Ruhr of the University Alliance Ruhr, Faculty of Chemistry, University of Duisburg-Essen, Essen, Germany; Centre of Water and Environmental Research (ZWU), University of Duisburg-Essen, Essen, Germany; Department of Soil and Crop Sciences, Colorado State University, Fort Collins, CO, USA; Energy and Environment Directorate, Pacific Northwest National Laboratory, Richland, WA, USA; Earth and Biological Sciences Directorate, Pacific Northwest National Laboratory, Richland, WA, USA; School of the Environment, Washington State University, Pullman, WA, USA; Centre of Medical Biotechnology (ZMB), University of Duisburg-Essen, Essen, Germany; DOE Joint Genome Institute, Lawrence Berkeley National Laboratory, Berkeley, CA, USA

**Keywords:** Stream, Microbial activity, Hyporheic zone, Anthropogenic stress, Microbiome, Climate change, Temperature

## Abstract

**Background:** Stream hyporheic zones represent a unique ecosystem at the interface of stream water and surrounding sediments, characterized by high heterogeneity and accelerated biogeochemical activity. These zones are increasingly impacted by anthropogenic stressors and environmental changes at a global scale, directly altering their microbiomes. Despite their importance, the current body of literature lacks a systematic understanding of active nitrogen and sulfur cycling across stream sediment and surface water microbiomes, particularly across geographic locations and in response to environmental stressors.

**Results:** Based on previously published and unpublished datasets, 363 stream metagenomes were combined to build a comprehensive MAG and gene database from stream sediments and surface water including a full-factorial mesocosm experiment which had been deployed to unravel microbial stress response. Metatranscriptomic data from 23 hyporheic sediment samples collected across North America revealed that microbial activity in sediments was distinct from the activity in surface water, contrasting similarly encoded metabolic potential across the two compartments. The expressed energy metabolism of the hyporheic zone was characterized by increased cycling of sulfur and nitrogen compounds, governed by *Nitrospirota* and *Desulfobacterota* lineages. While core metabolic functions like energy conservation were conserved across sediments, temperature and stream order change resulted in differential expression of stress response genes previously observed in mesocosm studies.

**Conclusions:** The hyporheic zone is a microbial hotspot in stream ecosystems, surpassing the activity of overlaying riverine surface waters. Metabolic activity in the form of sulfur and nitrogen cycling in hyporheic sediments is governed by multiple taxa interacting through metabolic handoffs. Despite the spatial heterogeneity of streams, the hyporheic sediment microbiome encodes and expresses conserved stress responses to anthropogenic stressors, *e.g.*, temperature, in streams of separate continents. The high number of uncharacterized differentially expressed genes as a response to tested stressors is a call-to-action to deepen the study of stream systems.

## Background

Streams connect habitats and ecosystems [1–3], scaling from small currents often originating in mountains, over mid-sized upland rivers to wide lowland streams feeding into the sea [4,5]. The community of microorganisms living within the stream are shaped by the distinct geomorphological and hydrodynamic features along the flow path [6,7]. In turn, stream microorganisms perform important ecosystem services, in surface water and underlying sediment [8,9]. A systematic approach to characterize streams is presented by the stream order metric also known as the Horton-Strahler number starting at 1 and increasing by the joining of streams with the same or lower order [10,11]. Headwater streams with low stream orders are often characterized by high variations of flow velocity, discharge, and temperature, and large amounts of terrestrial input like leaf litter [3,12,13]. Contrastingly, streams classified as higher stream orders are generally wider and faced with less variation compared to low order streams, carry more nutrients and shift towards an autotrophic system according to the River Continuum Concept (RCC) [3]. The change from headwater stream to the mouth is reflected in a change of fish communities [14,15], and both eukaryotic microbes [9] and prokaryotic microorganisms [16,17]. Corresponding changes in microbial community functions were shown to be related to the surrounding land use and cover, with differences for streams in wetlands, forests, and agriculture, respectively [16,18].

Apart from natural variability in environmental factors, anthropogenic influence in the form of stressors on streams also varies within the water column, both in type and intensity [19–22]. In Europe, only 37% of freshwater ecosystems are in a good or high ecological status including streams of all orders [23]. For example, artificial hydromorphological adaptations in the form of channels or dams are prominent stressors for streams at a global scale [24,25]. Anthropogenic pressures like inflow of wastewater can be concentrated around densely urbanized areas or human activities like mining can lead to inflow of nutrients and increased temperatures at industry sites [26,27]. By contrast, temperature increase due to human-caused climate change affects streams across all orders, potentially even more glacier-fed streams in mountain areas [28,29].

Natural and anthropogenic stress influence the ecosystem services provided by the organisms living in the surface water and sediment of streams [30,31]. However, an exhaustive study covering 1,851 sites in rivers and streams across North America found that healthy biological communities which are essential for provision of ecosystem services were only found in 28% of investigated stream miles between 2018 and 2019 [32]. In particular, the interface of surface water and groundwater, referred to as the hyporheic zone, represents a biological hotspot [33] that is easily prone to stressors [34,35]. Yet, microbial communities in the hyporheic zone are involved in complex nutrient cycling and actively shape the ecosystem structure [33,36–39]. They are not directly compositionally comparable to those of the overlying surface water column as demonstrated in, *e.g*., glacier-fed streams [40,41].

Multiple approaches can be utilized to study the effect of stressors and environmental variables on the interplay of stream microbial biodiversity and activity, such as the deployment of large-scale outdoor mesocosm experiments simulating stressors [42–44], or systematic field sampling of streams of different orders and habitat types [17,18,45,46]. The Worldwide Hydrobiogeochemistry Observation Network for Dynamic River Systems (WHONDRS) represents such a field study with 97 streams sampled in the summer of 2019 sampling study [47], leading to majority of sampling in the Genome Resolved Open Watersheds database (GROWdb) [18]. GROW revealed that the surface water microbiome of streams is characterized by aerobic and phototrophic metabolisms and follows the RCC by changing along the river gradient. However, this study also revealed the impact of environmental factors like stream order or land use on microbial assembly and activity [18]. Yet, apart from selected studies like GROW, genome-resolved activity data from metatranscriptomics analyses remains limited, as many studies either fail to recover metagenome-assembled genomes (MAGs), obscuring individual microbial contributions, or do not incorporate metatranscriptomics, masking transcriptional activities [16,17,48–50].

The full-factorial ExStream system which has been applied for organisms of multiple trophic levels in streams worldwide is an example of targeted stressor testing by manipulating abiotic factors [42,43,51–55]. In studies based on this mesocosm system and focusing on the microbiome of streams, the complex interaction of multiple anthropogenic stressors has been shown based on both metagenomic and metatranscriptomic analyses, especially for temperature and hydromorphological changes [30,54,55]. Both approaches, *i.e.,* field studies and mesocosms, have advantages and drawbacks. For example, field studies of multiple stream sites typically include a multitude of environmental variables which affect the microbiome and also enable extrapolation of obtained results to other catchments. Yet, the interplay of abiotic and biotic factors in natural systems may be too complicated for clearly linking community response to stressors. On the other hand, mesocosm studies offer the possibility to reduce complexity compared to field studies and increase statistical power due to replication. Yet, these advantages stand in contrast to limited generalizability by mesocosms located at single streams. Both approaches, field studies of multiple sites and replicated mesocosms, benefit from a high number of samples, whose combination helps to overcome low assembly quality of individual samples due to high complexity of microbial communities and to recover high-quality metagenome-assembled genomes (MAGs) [18,55–57].

Here, we overcome obstacles such as limited generalizability from single experiments and hampered recovery of microbial diversity from single samples, by combining hundreds of metagenomes from both field sampling and mesocosm experiments to generate two comprehensive river databases, one for MAGs and one for genes. By mapping metatranscriptomic and metagenomic reads obtained from 13 stream samples with data on both hyporheic sediment and surface water to our databases, we show that metabolic activity is not always consistent with encoded potential. For example, hyporheic sediments showed increased activity compared to the overlying surface water despite presenting similarly encoded metabolic potential. Additionally, we projected previously identified stress responses in mesocosm experiments to a diverse set of river sediment metatranscriptomes (n=23). This revealed a consistent active stress response across streams in North America.

## Methods

### Study design and aim

This study was designed with the aim of deciphering the relationship between microbial activity of hyporheic zone sediments and environmental factors like temperature change, while identifying and characterizing key microbial players governing biogeochemical cycling. To this end, we combined previously published [18,50,55] and unpublished data from 363 stream metagenomes building a comprehensive MAG and gene database to establish GROWdb version 2 (GROWdbv2, methods below). We coupled GROWdbv2 to sediment (n=23) and surface water metatranscriptomic data (n=13) to identify distinct metabolic patterns in these two connected ecosystems. Additionally, sediment metatranscriptomes were used to infer the response of the microbiome to the environmental factors by testing differential gene expression (DGE) for temperature (8.5 to 31.4 °C at sampling), sediment composition (16.7% to 94.1% total sand), and stream orders (1 to 8), comparing the response derived from microcosm experiments in ExStream which clearly constrained stress responses of individual biomes [30,54,55,58].

### Collection of river sediment and surface water samples

Sampling and processing of sediment and surface water of global rivers samples has been conducted within the project Worldwide Hydrobiogeochemistry Observation Network for Dynamic River Systems (WHONDRS, https://www.pnnl.gov/projects/WHONDRS) [47], with surface water data previously reported [18]. Sampling followed the protocol for the summer 2019 sampling study [59]. Briefly, sediment was taken within a 1 m^2^ area from the streambed (1-3 cm depth) at five to ten locations and preserved in RNALater. Surface water (approximately 1 L) was filtered through 0.33 μm sterivex filters (EMD Millipore), which were capped and filled with 3 mL RNALater. Samples were stored on wet/blue ice or in a 4 °C refrigerator until shipped to Pacific Northwest National Laboratory (PNNL, USA), where samples were subsampled and frozen at −20 °C.

### Nucleic acid extraction and sequencing

Surface water DNA extraction and sequencing methods were previously reported [18]. Here, we build upon the previous study by incorporating the paired sediment metagenomic and metatranscriptomic data. Coextraction of DNA and RNA from sediment was done at Colorado State University (Fort Collins, USA) using NucleoBond RNA Soil kit and DNA Set for NucleoBond RNA soil kit coupled with RNA Clean & Concentrator-5 (Zymo Research Cat. # R1013). Metagenomic and metatranscriptomic sequencing was performed at the Joint Genome Institute (JGI) under the umbrella of a Community Science Program (CSP; proposal: 10.46936/10.25585/60001289; award: 505780). Metagenomic sequencing followed the protocol described in [18]. Briefly, DNA fragmentation and adapter ligation was done using the Nextera XT kit (Illumina) and unique 8 bp dual-index adapters (IDT, custom design). After enrichment, prepared libraries were sequenced on an Illumina NovaSeq sequencer according to a 2 × 150 nucleotide indexed run program. Metatranscriptomic sequencing and library prep was done using JGI established protocols based on rRNA removal (Qiagen FastSelect probe sets for bacterial, yeast, and plant rRNA depletion (Qiagen)), Illumina TruSeq Stranded mRNA Library prep kit (Illumina), and Illumina NovaSeq sequencer following a 2×150 nucleotide indexed run recipe. Collectively, in this study we publish 53 metagenomes and 23 metatranscriptomes from sediment, analyzing them together with previously published 310 metagenomic datasets [18,41,50,55].

### Metagenomic data processing and construction of GROWdbv2 MAGs

We combined multiple existing datasets with the aim of compiling a comprehensive database of metagenome-assembled genomes (MAGs) from river ecosystems around the world. From the new metagenomes (n=53) reported here, we recovered 1,696 new MAGs following protocols established by GROWdb [18]. Briefly, metagenomic reads were trimmed using sickle (v1.33, [60]) and assembled with IDBA-UD (v1.1.0, [61]) or MEGAHIT (v1.2.9,[62]) and resulting contigs subsequently binned with metabat2 (v2.12.1, [63]). In total, our database comprised 6,724 MAGs originating from GROWdb (3,824 previously published MAGs [18] and 1,696 unpublished MAGs), *ExStream* mesocosm experiment (69 MAGs, [55]), Erpe river study (1,033 MAGs, [41]), and Columbia river study (102 MAGs, [50]). Taken together, 5,915 originated from surface water samples, 478 from sediment, and 331 from porewater of sediments. A dereplicated set of medium and high-quality MAGs with >50% completion and <10% contamination was obtained by running dRep (v3.5.0, [64]) at 99% identity with CheckM2 quality assessment as input (v1.0.2, [65]).

The resulting 3,741 MAGs, representing GROWdbv2, were taxonomically annotated using the classify_wf workflow of GTDB-Tk (v2.4.0, [66]) with reference data version r220. The concatenated sequence of all genomes was used for gene prediction using prodigal (v2.6.3, [67], -q -f gff -m -p meta) and a mapping index was built using Bowtie2 (v2.4.5, [68]). Metabolic annotation was performed with DRAM (v2beta, [69]) based on predicted genes using the following modules: CAMPER, dbCAN, Genome_stats, KEGG, MEROPS Peptidases, Heme Regulatory Motifs Counts, and Pfam. Genes playing a part in microbial energy metabolism were grouped using the distill sheets (distill_energy_Jan252024.tsv) as part of DRAM.

### Metatranscriptomic data processing and microbial expression calculation of MAGs

Metatranscriptomic sequences were quality checked and trimmed following the workflow established by [18]. Briefly, trimming filtering was done using bbduk and rqcfilter2 embedded in BBTools (Bushnell, https://jgi.doe.gov/data-and-tools/bbtools/bb-tools-user-guide/). The microbial fraction of metatranscriptomic reads was estimated using SingleM (v0.18.3, [70]).To assess MAG activity, trimmed and quality-checked metatranscriptomic reads were mapped using Bowtie2 (v2.4.5, -D 10 -R 2 -N 1 -L 22 -i S,0,2.50, [68]), filtered at 97% minimal identity and used to calculate counts of predicted genes with htseq-count (v2.0.5, [71]). Counts were transformed to geTMM (gene length corrected trimmed mean of M-values) using edgeR package [72].

### Identification and characterization of MAGs active in sediments

Lineage-specific microbial activity in sediments was obtained by investigating MAGs active in those samples. For that, genes were only considered if expressed in at least 5% of samples and a MAG was counted as active with a minimum of 20 actively expressed genes following [18]. Sulfur and nitrogen pathways encoded and expressed in MAGs based on annotations by DRAM were visualized using Affinity Designer (v1.10.8). A phylogenetic tree of active bacterial MAGs was inferred via GTDB-Tk (v2.4.0, [66]) based on the de_novo workflow with “p Acidobacteriota” as outgroup. Plotting was done in R using packages ggtreeExtra [73] and ggnewscale [74]. In case of homologous genes, *i.e., nxr* vs. *nar* and *amo* vs. *mmo*, identity of genes was verified using phylogenetic analyses. First, sequences were aligned using mafft (v7.407, -auto, [75]). Then, the alignment was trimmed using trimAL (v1.5.rev0, -automated1, [76]) and a tree was built with IQ-TREE (1.6.12, -alrt 1000 -bb 1000 -m C20+G+F, [77]). Accession numbers of reference genes, partially compiled from [78], used herein can be found in **Figures S2** and **S3**.

### Clustering, annotation, and mapping of genes from metagenomic assemblies

A gene database was built from assemblies of 363 metagenomes corresponding to those used for the MAG database. All contigs/scaffolds from metagenomic assemblies were filtered to a minimal contig/scaffold length of 1500 base pairs and concatenated. Gene prediction, annotation, mapping of quality-checked metatranscriptomic and metagenomic reads (trimming done as described in [18]), and counting of hits was done as explained above for the MAG analysis. Proteins were clustered using MMseqs2 at 95 % amino acid identity [79] and singletons were excluded from further analyses resulting in 11,043,548 clusters. To enhance read recruitment, reads were mapped against all genes for all assemblies. Since this mapping did not solely contain clustered representative sequences, counts were summed by cluster after mapping. This procedure was done for all metatranscriptome samples and metagenomic samples.

### Effect of environmental factors on sediment gene expression

Transformation to geTMMs for resulting gene expression counts was done as explained above with the same thresholds. Based on the 23 metatranscriptomic sediment samples, the influence of abiotic environmental factors on the microbial activity was assessed. Specifically, temperature, water depth, percentage of total sand in sediment, nitrogen and carbon content, and stream order were extracted for the respective sampling points from WHONDRS [59,80]. Stream order was included in this study as a factor as it has been proven to be a significant factor for the microbiome assembly of stream surface water [18] and is a known metric to compare streams across catchments. Multivariate statistics were done based on the Bray-Curtis Dissimilarity on normalized counts. These included permanova tests (adonis2, permutations=999, by=”margin”) and nonmetric multidimensional scalings (NMDS) [81–84]. The richness of active genes per sample and the number of shared active gene clusters between samples was visualized using UpSetR [85]. The count of active genes per sample was correlated using the native R stats package with the prokaryotic fraction in metatranscriptomic reads as inferred by SingleM (v0.18.3, [70]). Likewise, prokaryotic fraction in metatranscriptomic reads was also correlated with the temperature of the sample.

Differential gene expression was tested for the three most significant environmental factors as inferred by multivariate statistics, *i.e.,* temperature, percentage of total sand, and stream order. Therefore, the dispersion of gene counts was estimated using estimateDisp function (design=Temperature+Percent_Tot_Sand+Stream_order, robust=T) and a generalized linear model fitted with glmQLFit embedded in edgeR (v4.0.16; [72]). Significant up- and downregulated genes were identified with an adjusted p-value < 0.05 (Benjamin-Hochberg) and an absolute log2-fold change of 0.75. For the respective genes, functions were annotated using DIAMOND blast (v2.0.15; blastp --fast -e 0.00001 -k 1; [86]) against FunTaxDB (v1.4 from Nov. 2023; [87]), based on UniRef100 [88]. Data was plotted using ggplot2 [89].

## Results

To obtain a comprehensive overview of metabolic activity in stream sediments as a function of environmental factors, 23 samples from 20 different streams were surveyed. Environmental factors measured across all sites included surface water temperature that ranged from 8.5 °C to 31.4 °C at the time of sampling, sediment sand composition that varied between 16.7% to 94.1% total sand, and stream orders from 1 to 8 (**Figure 1**).

**Figure 1:**
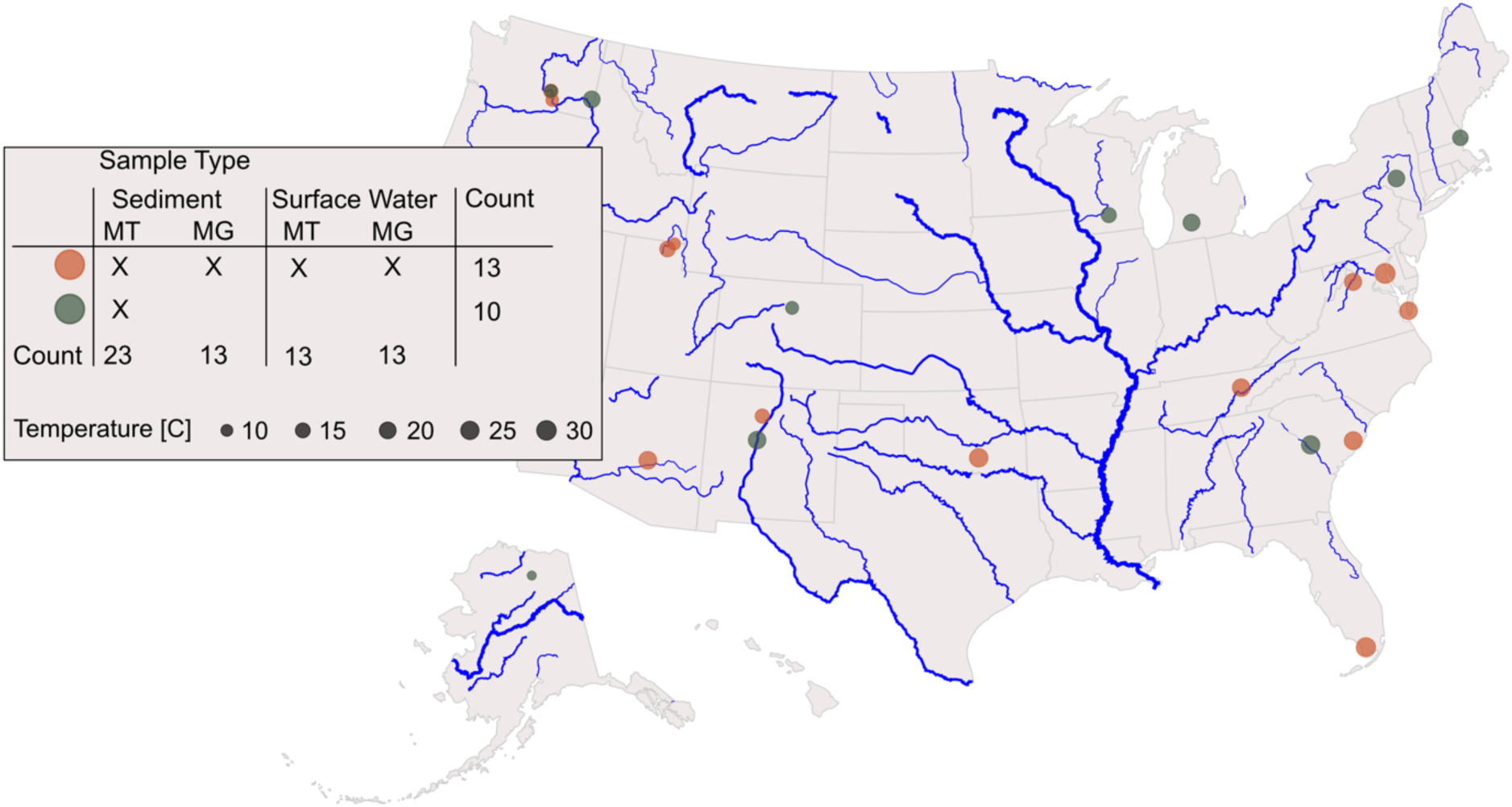
Geographic locations of river samples across North America used in this study. In total, there were 23 samples, 13 of which have metagenomics and metatranscriptomics data from both sediment and the associated water column (orange). Ten of the samples were sediment metatranscriptomes only (green).

Metatranscriptomes and metagenomes were mapped to two comprehensive databases built for this study. First, 6,724 metagenome assembled genomes (MAGs) were dereplicated at 99% identity to 3,741 medium and high-quality genome representatives forming the baseline for genome-centered analysis. The built gene database contained 11,043,548 clusters (95% AAI) and was used for detailed analysis of stream metabolic functions and microbiomes’ response to environmental stressors.

### Metabolic activity of stream sediments is greater than that of surface water

Genetic information of all microbiomes across the 13 samples encoded for main energy conservation pathways in both sediment and surface water, as per the comprehensive river gene database built in this study (**Figure 2**). While surface water encoded significantly more genes whose products are involved in sulfur and methane cycling, more enzymes for nitrogen and other C1 cycling were encoded in sediment metagenomes (Kruskal-Wallis-Test, Benjamini-Hochberg-adj. p-value < 0.05, see **Figure 2**). This finding is in stark contrast with gene expression measured via transcriptomics in this study, highlighting the necessity to include metatranscriptomics in river ecosystem exploration. As expected, not all genes of all pathways recruited reads from metatranscriptomes as certain genes might be expressed in too small a number to be captured in metatranscriptomics (if expressed at all). For example, “anoxic photosystem II” was encoded in all but one sample, but only detected in metatranscriptomes of three out of 13 surface water samples and six out of 13 sediment samples (**Figure 2**). Besides the discrepancy between functional potential and measured gene expression, significantly higher expression profiles (Kruskal-Wallis-Test, Benjamini-Hochberg-adj. p-value < 0.05, see **Figure 2**) were detected in the sediment samples compared to the surface water. Most striking, pathways that were more frequently encoded in the surface water like methanogenesis (including, *e.g*., key gene, *mcrA*, for methanogenesis), sulfur cycling (including, *e.g*., key gene, *dsrA/B*, for dissimilatory sulfate reduction), or hydrogen oxidation/production displayed significantly higher expression patterns in the sediment samples (Kruskal-Wallis-Test, p-value < 0.05, see **Figure 2**). Notably, the two sediment samples with the highest temperature during sampling, *i.e.*, shark river slough (“sharkriverslough_0042”, Florida, USA) and muddy creek (“muddycreek_0082”, Maryland, USA), exhibited consistently fewer expressed pathways than those in other sediment samples, particularly pathways related to methane metabolism. This deviation to the other sediment samples is a first indicator for a temperature dependency of the microbial activity within sediments. If these two samples had exhibited expression profiles more similar to the other sediment samples, the difference between surface water and sediment might have appeared even more pronounced, with more pathways showing significantly higher expression.

**Figure 2:**
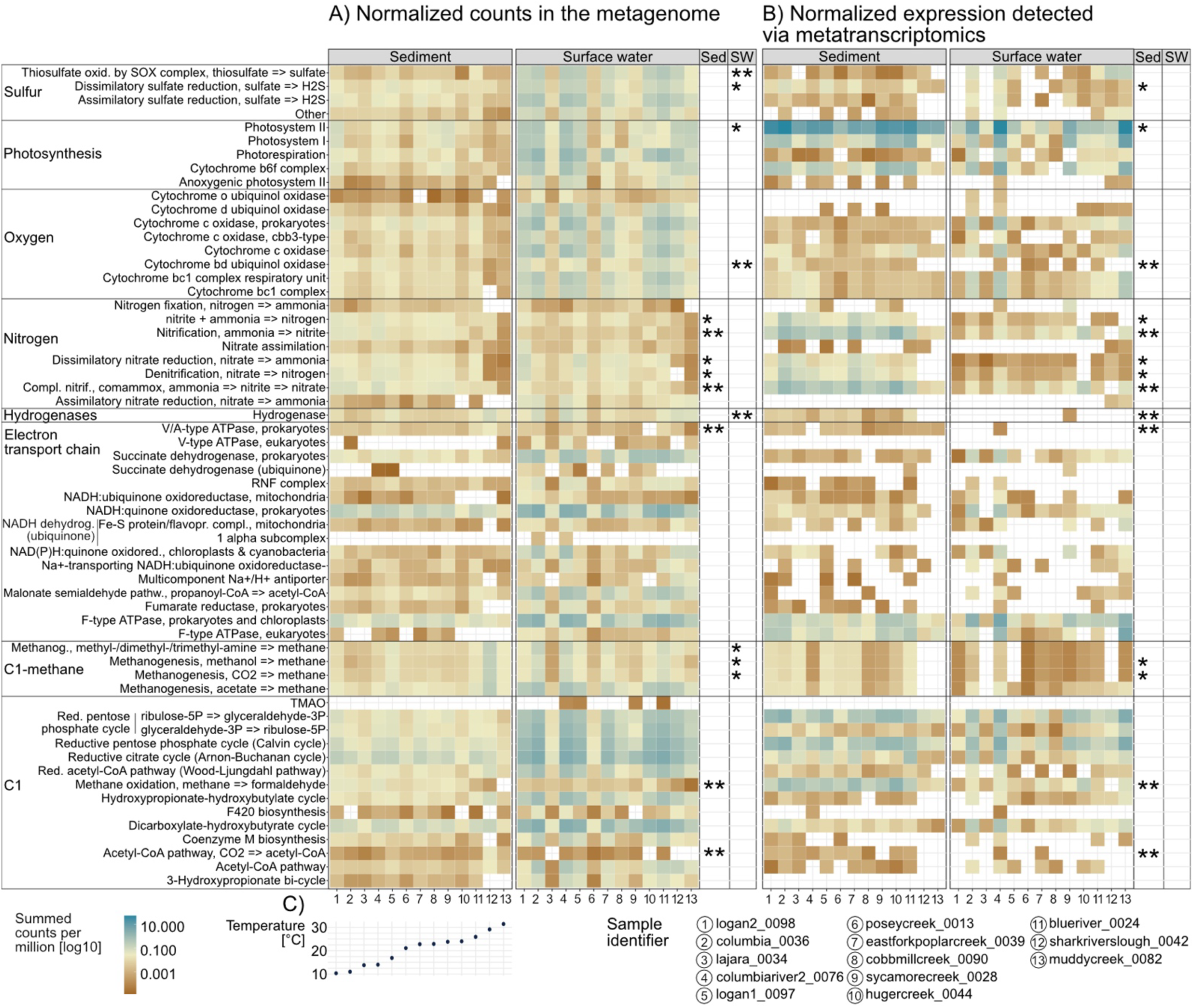
Comparison of microbial energy metabolism in sediment and surface water, encoded in the metagenome (A), and its activity measured via metatranscriptomics (B). Samples were sorted according to increasing temperature (C). The mean counts per pathway were tested for significant differences between surface water and sediment per method (Kruskal-Wallis-Test, n=13, visualized in Fig. S1); a significantly higher mean is marked by significance indices in the respective column (* and ** for adj. p-value (BH) < 0.05 and < 0.01, respectively). While in the metagenome selected functions are more abundant in surface waters than sediment samples, expression rates are always greater in the sediment samples compared to surface water, if significant differences were detectable.

### Sulfur and nitrogen cycling in stream sediments is a community endeavor

The analysis of metagenome-assembled genomes (MAGs) based on our river MAG database followed by expression analysis across thirteen samples (**Figure 1**), was in agreement with transcriptomics analyses for which we used protein clusters (**Figure 2**), *i.e*., mostly organisms connected to sulfur and nitrogen cycling were active in the sediment (**Figure 3**). Thus, the MAG-centered analysis was framed for the hyporheic sediment with 75 MAGs spanning ten bacterial phyla identified as active (**Figure 3A**). For example, MAGs belonging to phylum *Nitrospirota* (MAG identifier 13-16) were found to be active by showing gene expression only in sediments and not in the overlying surface water.

**Figure 3:**
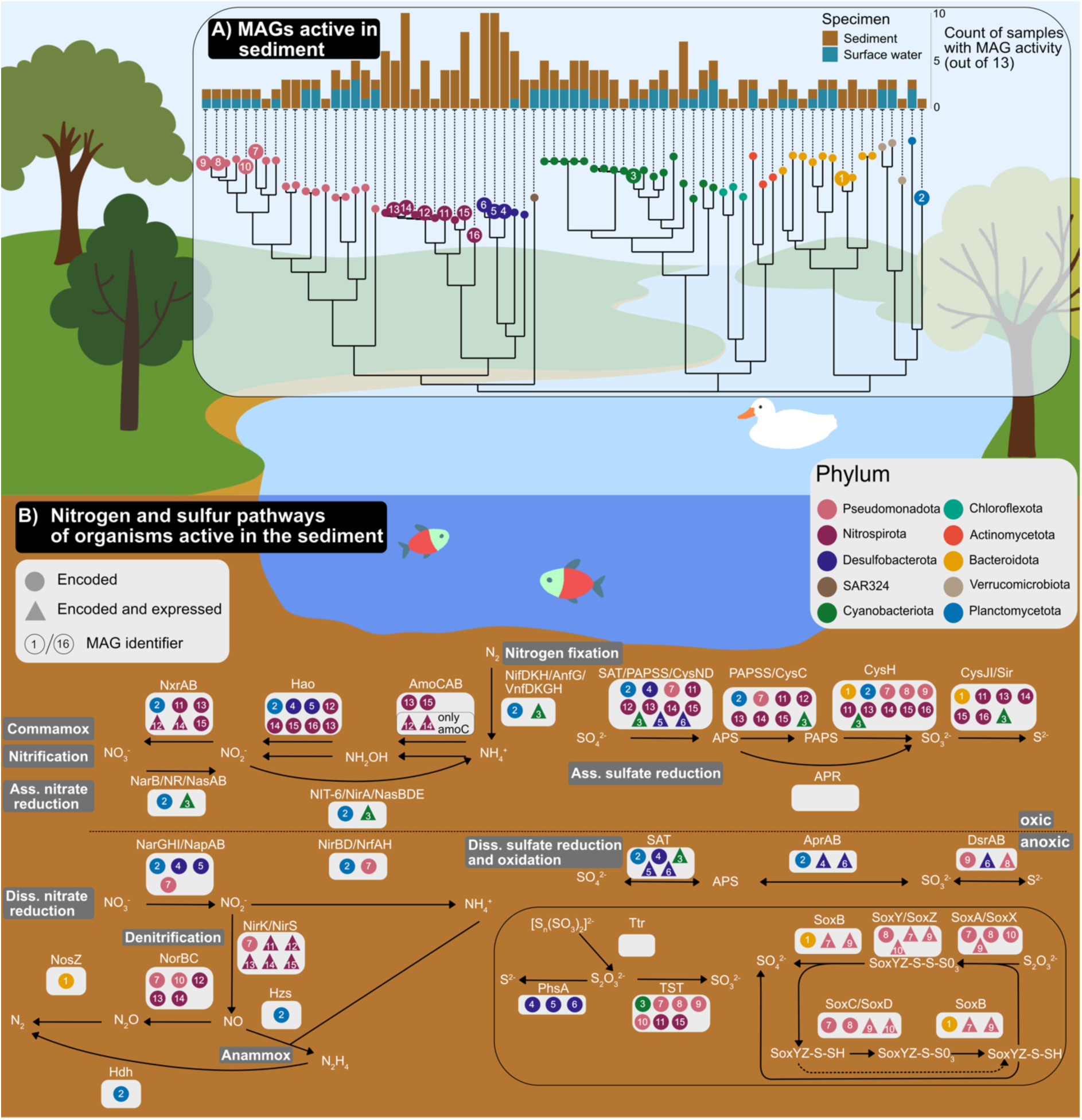
Schematic overview of key metabolic processes governed by MAGs active in the sediment. (A) Phylogenetic tree of MAGs found active in sediments (>=20 genes expressed) and number of samples with activity of respective MAGs in sediment and water samples, respectively. The MAG identifiers refer to genomes of organisms active in nitrogen or sulfur cycling (B). Genes of these pathways were expressed by organisms which were mostly found to be active in sediments only, especially Nitrospirota and Desulfobacterota. To be assigned as expressed by an organism, genes had to be expressed in at least one sediment sample. In case of reactions involving multiple enzymes, ⅔ of the respective genes had to be encoded or expressed to be included in the figure. For homologues such as amoA/B/C and mmoA/B/C and nxrA/B and narG/H, phylogenetic trees were inferred to assign the correct annotation (Figure S2 and S3).

A detailed reconstruction of the underlying pathways for both sulfur and nitrogen turnover in sediments revealed the interplay of multiple taxa (represented by MAGs) responsible for the two nutrient cycles. The close phylogenetic relationship between Nitrospirota and *Desulfobacterota* (**Figure 3A**) was also reflected by shared possession of genes associated with complete ammonia oxidation (COMMAMOX) and assimilatory sulfate reduction, respectively. Three Nitrospirota MAGs (MAGs 13-15) affiliated to family *Nitrospiraceae* (one of them genus *Palsa* (MAG 15)) encoded all genes for both pathways that are active under oxic conditions. Key enzymes of the COMMAMOX pathway, *i.e.*, *nxr and amo*, were also expressed by *Nitrospirota* (represented by MAGs 12 and 14). The metabolic versatility of this phylum was also demonstrated by expression of key denitrification genes like *nirK* that encodes for nitrite reductase which catalyses the reduction of nitrite to nitric oxide. Sulfur cycling by *Desulfobacterota* was likely mediated via dissimilatory sulfate reduction in the anoxic part of the sediment. Pseudomonadota encoded and expressed key pathways like the *Sox* system for sulfur oxidation.

Apart from these two main phyla being responsible for sulfur and nitrogen cycling, two other taxa were active across multiple pathways. As such, the Cyanobacteriota MAG (MAG 3) of family *Nostocaceae* encoded for active nitrogen fixation, assimilatory nitrate reduction, and assimilatory sulfate reduction across samples. Another generalist for sulfur and nitrogen cycling in stream sediments was represented by a MAG annotated as *Candidatus Brocadia* sp, phylum Planctomycetota. This MAG encoded for most of the genes of sulfur and nitrogen cycling depicted in **Figure 3**, even for both homologues of the nitrate reductase enzyme (*narGH*) and nitrite oxidoreductase (*nxrAB*) (see **Figure S3**). Taken together, transcriptomes mapped to our MAG database demonstrated that multiple organsims like *Nitrospirota* were involved in nutrient cycling and deemed specialists as focused on specific pathways. We also identified multiple organisms that were rather generalists with high metabolic versatility and expressed functions that catalyzed both nitrogen and sulfur cycling.

### Microbial activity in the hyporheic zone is influenced by temperature and sediment composition

Although we detected explicit patterns between sediment and water samples, microbial gene expression was not detected to be homogeneous across all sediment samples (**Figure 2**), suggesting a meaningful influence of environmental factors in the hyporheic zone. To test this hypothesis, key factors (*e.g.*, being important for streams ubiquitously and/or describe general aspects of streams) were selected from the accompanying WHONDRS metadata and used for statistical testing based on the expression profiles across all 23 sediment metatranscriptomes (**Table S1**). To comprehensively test for differential gene expression, we compiled an overarching river gene database from 363 samples comprising 11,043,548 representative genes, whose proteins were clustered at 95% amino acid identity. Temperature and percentage of total sand (differentiated from clay and silt) as continuous variables showed significant association with microbial activity (adonis2, p-value < 0.05, **Table S1**). Bray-Curtis dissimilarity-based nonmetric multidimensional scaling (NMDS) revealed a clear shift of the microbial expression profile with temperature increase (**Figure 4a**), which was accompanied by lower gene expression in samples with higher temperatures (**Figure 4b**). Correlating the prokaryotic fraction of the microbial activity, as estimated based on metatranscriptomic reads using SingleM [70], with temperature supported the hypothesis that the observed decrease of microbial activity at higher temperatures might be due to an increase of eukaryotic activity (**Figure 4c**). An increase of eukaryotic abundance is, in turn, suggested to be negatively correlated with the calculated prokaryotic fraction. Samples with the highest total count of expressed genes shared the most genes (1,043 genes) with each other. All samples shared 96 actively expressed genes (**Figure 4b**), of which 20 were annotated as Photosystem I/II and 40 as uncharacterized.

**Figure 4:**
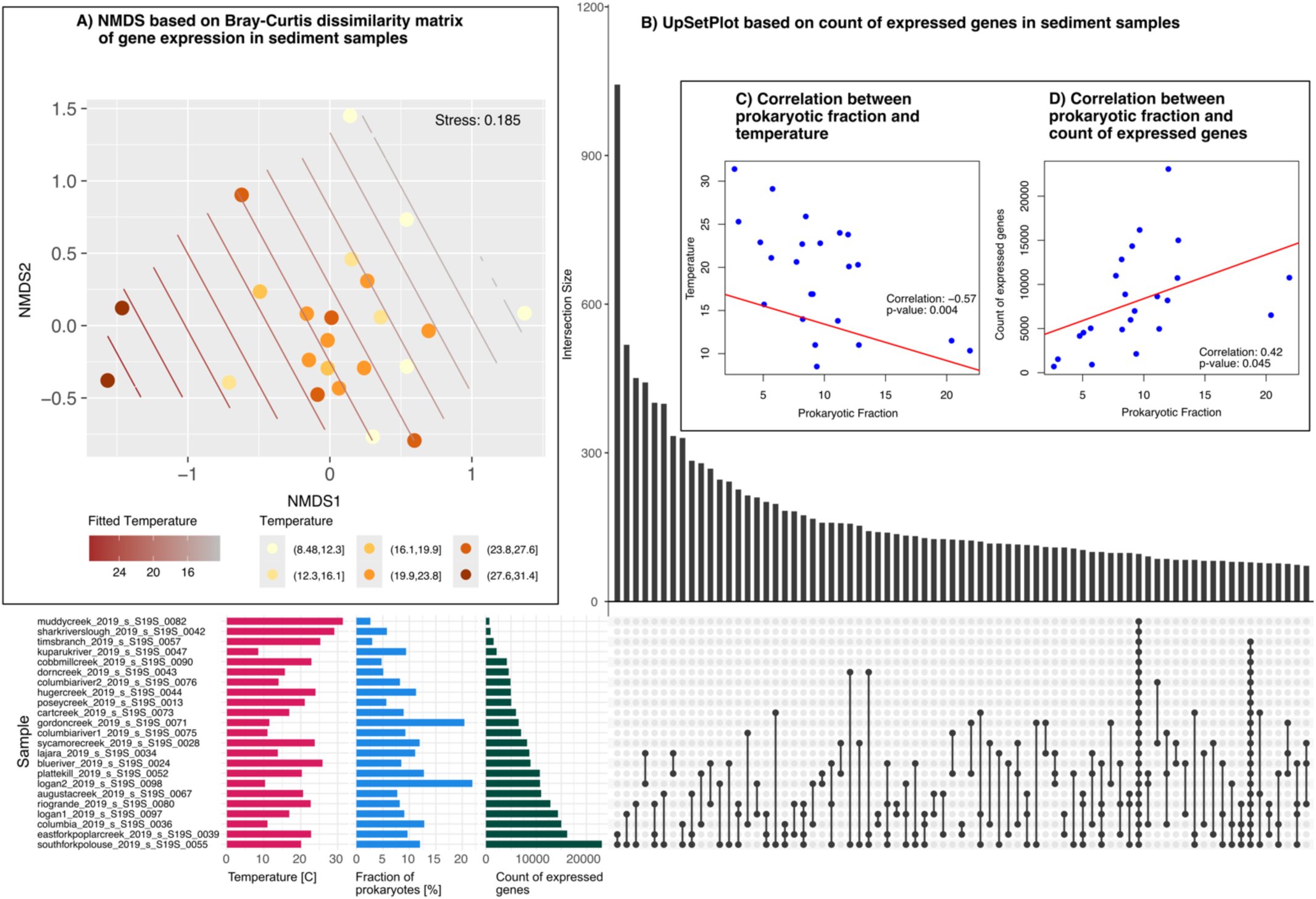
Influence of environmental factors on gene expression in stream sediments. A) Temperature effect on gene expression depicted by NMDS with fitted temperature and grouping of samples into six temperature intervals. B) UpSetPlot of active genes shared between samples (only top 75 interactions shown). Samples with the highest temperature have the least expressed genes. C) and D) Correlation between prokaryotic fraction and temperature and count of expressed genes, respectively. The prokaryotic fraction of metatranscriptomic reads was inferred with SingleM [70]. There was a significant negative correlation with temperature, and a positive with the count of active genes.

### Temperature activates conserved microbial stress response across stream sediments

To infer the specific response of the microbiome to environmental factors, differential gene expression based on the three factors that were identified to have the most significant effect on community gene expression (**Table S1**), *i.e.,* temperature, percentage of total sand, and stream order (**Figure 5**) was performed. Further studies with additional replication would include other factors like depth or nitrogen/carbon percentage.

**Figure 5:**
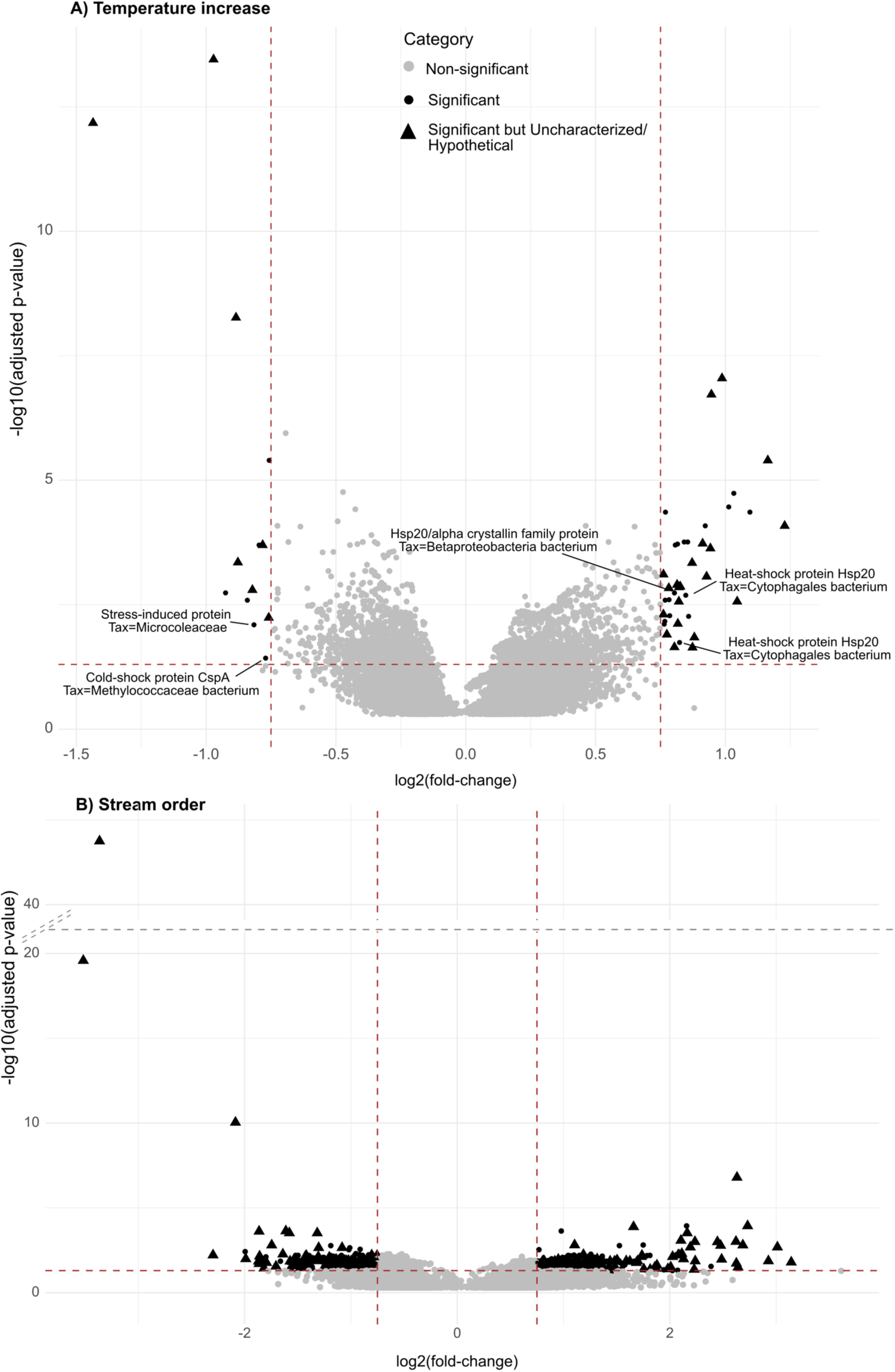
Differential gene expression (adj. p-value < 0.05 and abs(LFC) > 0.75) due to temperature and stream order. A) Temperature increase caused significant upregulation of heat-shock proteins while cold-shock proteins were downregulated (52 differentially expressed genes in total). Annotation was only shown for genes directly related to temperature or stress response. B) Stream order caused significant up and downregulation of a large set of in total 1,098 genes. Top differentially expressed genes with abs(LFC) > 1 are shown in Supplementary Figures S4 and S5.

Significant differential expression (adj. p-value < 0.05 and abs(LFC) > 0.75) was detected for temperature (**Figure 5a**) and stream order (**Figure 5b**), but not for percentage of total sand which is contrary to results from PERMANOVA testing (**Table S1**). With increasing temperature, stress responses like heat-shock proteins (*e.g.*, Hsp 20), as well as energy-related genes like Photosystem I/II were upregulated. By contrast, significant downregulation was detected for cold-shock protein CspA and a gene annotated as “stress-induced protein”. Compared to the small number of 53 differentially expressed genes by temperature, stream order change led to 1,098 differentially expressed genes, of which 416 up- and 734 downregulated genes were functionally annotated (**Figure 5b**). Upregulated genes at higher stream order included seven bacterial genes annotated as heat-shock proteins (Hsp20), other stress response genes like the anti-sigma factor antagonist gene, seven genes encoding cold-shock proteins, or multiple genes regulating translation and transcription or annotated as transposases. Prominent downregulation at high stream order was found by 22 cold-shock proteins. Overall, stress responses were actively transcribed at low stream order and included anti-sigma factors, chaperones, three heat-shock proteins, histones, response regulators, or transcription/translation genes (**Figure 5b**).

Focussing on the taxonomic annotation of differentially expressed genes associated with stream order change, 173 belonged to *Nitrospirota*, of which 163 were downregulated with increasing in stream order. Downregulated genes included ribosomal proteins and nitrate oxidoreductase subunits. While downregulated photosystems were annotated with typical freshwater Cyanobacteria, mainly *Chamaesiphon* sp., upregulated photosystems were linked with species also found in marine environments like *Thalassiosira nordenskioeldii.* Similarly, temperature increase also had an effect on gene expression by Cyanobacteria in the form of seven upregulated genes that were taxonomically annotated to genera *Planktothricoides* sp. and one as *Planktotrix*.

In total, 854 functionally annotated genes were found to be differentially expressed based on changes in either temperature or stream order. A major fraction of these representative genes (216) were identified in the included mesocosm experiment with 64 different metagenomes generated in Germany [55] reflecting that (a) the applied mesocosm approach is valid to study stressor effects in streams and (b) found stressor effects from one continent are transferable to a wide range of catchments from another continent.

## Discussion

Streams represent unique ecosystems with a high spatial heterogeneity, not only with respect to the influence of abiotic factors like surrounding land usage and cover, but also to the resulting microbial community living within the surface water [16,18] and within the sediment and with the hyporheic zone as a hotspot of activity [17,33]. While already shown for glacier-fed streams [40], we herein establish that the two habitats –surface water and hyporheic zone– exhibit distinct gene expression profiles for multiple key metabolic pathways even across different stream orders. Increased expression of genes encoding key metabolic pathways in sediment samples including those with lower than average prokaryotic activity agrees with the concept that describes the hyporheic zone of streams as the ‘river’s liver’ [37]. In addition, our findings suggest that there are conserved metabolic capacities in sediments spanning all stream orders. Highly active nitrogen and sulfur metabolisms within the sediment microbiomes may also be a microbial response to high levels of anthropogenic land use that introduce such nutrients in the form of fertilizers and storm water runoff (*e.g*., stream Logan, Utah, with sample “logan2_0098”) or wastewater treatment effluents (*e.g.*, stream East Fork Poplar Creek, Tennessee, with sample “eastforkpoplarcreek_0039”) [90–92].

A comprehensive river genome catalogue compiled from worldwide river metagenomic studies [18,47,50,55] including 3,741 genomes enabled identification of *Nitrospirota* and *Desulfobacterota* as main players in nutrient cycling in stream sediments. Similar to lake ecosystems [93], cycling of sulfur and nitrogen compounds seems to be a community effort shared by multiple lineages. Such interactions between microbial organisms has previously been described as “metabolic handoffs” for groundwater ecosystems [49]. This concept can now be extended to river ecosystems, rendering this a key concept for freshwater sediment microbiomes. One example of a metabolic handoff identified in the hyporheic zone is partial denitrification with *nirK*, found active only in *Nitrospira* sp.. In contrast, other steps of denitrification were found to be encoded in organisms belonging to *Desulfobacterota*, *Pseudomonadota*, and *Planctomycetota* in case of *narGHI/napAB,* and *Bacteroidota* in case of *nosZ*. While the role of *nirK* in Nitrospira, whose organisms participate in the nitrogen cycle mainly via complete ammonia oxidation (COMMAMOX, [94,95]), is still not completely understood [96], this finding suggests the metabolic potential for reduction of NO_2_^−^ with other organic substrates as electron donors [97] in river sediments. Elimination of NO_2_^−^ in freshwater is an ecologically important mechanism with, *e.g.*, increasing NO_2_^−^ concentrations during sulfur-based denitrification thereby constraining nitrogen cycling [98,99]. Beyond nitrite reduction, the high metabolic versatility of sediment *Nitrospirota* also includes assimilatory sulfate reduction encoded in multiple genomes in this study, which has previously been discussed as an alternative electron and energy source in one Nitrospira species, *i.e.*, *N. gracilis* [100]. These results are a call-to-action to further investigate Nitrospira in rivers along with members of Planctomycetes, as one organism whose MAG was annotated as *Ca*. Brocardia sp. also showed high metabolic versatility based on the encoded genes. Taken together, active genes in the anoxic and oxic metabolic pathways of the river sediments suggest activity of organisms that can swiftly adapt to changing conditions of the hyporheic zone and its available nutrients, especially in the upper 5-cm sediment layer.

Our finding that water temperature is a main factor shaping microbial activity in the hyporheic zone is in agreement with results from previous mesocosm experiments [55]. As such, we can verify the applicability of mesocosm experiments like *ExStream* [42,43,51–55] also for microbes of the hyporheic zone. Elevated stream water temperature has previously been related to an increase of eukaryotic growth in the form of algal blooms [101,102], which explains our correlation of fewer gene transcripts with decreases in prokaryotic signals. Comparing our findings with those from mesocosms in Germany [55], we identified a uniform upregulation of temperature-related stress response (*e.g.*, heat-shock protein Hsp20) for microbial communities living in the hyporheic zone across catchments of distinct continents. This uniform stress response suggests that organisms encoding such stress responses have an advantage at the cost of higher energy demand in contrast to more sensitive species during ecosystem degradation caused by anthropogenic stressors. The Asymmetric Response Concept [103], a conceptual framework for stressor effects on stream communities, postulates that species tolerances against stressors are of higher importance in phases of stress impact compared to phases of recovery. While this asymmetry has already been tested and partially verified by mesocosm experiments [55], this study extends the importance of stressor tolerance encoded and expressed by microbes to natural stream ecosystems.

It has been shown that the matrix of the hyporheic zone can influence the community composition of macroinvertebrates [104]. Based on our results from multivariate statistics this can now also be extended to microbial activity at the community level. Sediment composition is an example for the overall high spatial heterogeneity of the hyporheic zone, as its characteristics (*e.g.,* sand versus silt) influence the water flow and water chemistry in the sediment [105]. Being influenced by the sediment composition, hyporheic water flow can in turn affect the surface water temperature on small spatial scales below 0.25 m^2^ [106]. A detailed insight into how sediment composition and microbial activity are entangled was not obtained by this study based on differential expression analysis. Therefore, we suggest designing a targeted sampling strategy with replicates and different sampling points along a gradient of sediment composition within a single catchment and comparison across catchments.

The River Continuum Concept (RCC) aims to describe the river ecosystem and its longitudinal connectivity based on physical and biological gradients and has been shown to hold true for the microbial activity in the surface water [3,18]. With this study, we can extend this concept to the hyporheic zone. In detail, the high number of upregulated genes at low stream order may be related to higher variations of temperature, flow velocity, and nutrients in lower stream orders compared to wide rivers of high stream order [3,12,13]. In a multifactorial mesocosm study based on the *ExStream* setup [42,43,51–55], chaperones were found to be significantly upregulated due to lowered flow velocity [55]. Herein, we show that chaperones were along with other stress responses also related to stream order change. The detected higher expression rate of chaperons in low stream order samples suggests that the microbiomes need to adapt quickly to the rapidly changing environmental conditions of headwater streams compared to more stable high order streams. Thus, it could be hypothesized that anthropogenically induced changes of stream flow velocity could pose a similar stress on the microbes living in natural stream hyporheic zones as found in experimental setups.

Taken together, our results warrant further research investigating hydrologically connected stream networks, from headwater to sea to systematically expand the findings of our study. We would like to emphasize that this study benefited greatly from marrying datasets of different working groups and methodological approaches (field and mesocosm studies). Thus, this study reflects the strength of data sharing based on metadata standards and principles like the Data Reuse Information tag (DRI) [107] and FAIR principles [108]. Systematic follow-up studies which include datasets of previous stream research would also provide the opportunity to further characterize the high number of so-far unannotated genes in stream sediments, which harbor a yet undiscovered microbial diversity urging detailed biochemical and bioinformatic investigation.

## Conclusion

The hyporheic zone of stream sediments represents a unique ecosystem characterized by high spatial heterogeneity and high microbial diversity. By comparing 23 stream samples on the basis of comprehensive MAG and gene databases, we revealed an active core microbial community interacting in cycling of essential nutrients like nitrogen and sulfur compounds. At the community level, microbial activity was shown to be affected by sediment composition, temperature, and stream order with a conserved response to stress in the form of up and downregulation of specific genes. Our findings reveal that North American stream microbiomes are in a constant state of stress despite the unique physical and chemical characteristics of each catchment.

## Declarations

### Ethics approval and consent to participate

Not applicable.

### Consent for publication

Not applicable.

### Availability of data and material

All metadata and methods regarding sampling procedure are publicly available on the ESS-DIVE data repository from WHONDRS [47,59].

### Competing interests

The authors declare that they have no competing interests.

### Funding

This study was performed in context of the Collaborative Research Center (CRC) RESIST and analyses were done in Project A01, funded by the German Research Foundation (DFG) CRC 1439/1; project number 426547801. Tom L. Stach was supported by the German Academic Scholarship Foundation. Open Access funding enabled and organized by Projekt DEAL. WHONDRS efforts described here along with J.C.S. and A.E.G. were funded under the River Corridor Science Focus Area (RC-SFA) at Pacific Northwest National Laboratory (PNNL) and funded by the US Department of Energy (DOE) Office of Biological and Environmental Research (BER) Environmental System Science (ESS) Program. PNNL is operated by Battelle Memorial Institute for the DOE under contract no. DE-AC05-76RL01830. Metagenomic and metatranscriptomic sequencing was performed at the JGI under a Community Science Program. The work (proposal 10.46936/10.25585/60001289) conducted by the US DOE JGI (https://ror.org/04xm1d337), a DOE Office of Science User Facility, is supported by the Office of Science of the US DOE operated under contract no. DE-AC02-05CH11231.

### Authors’ contributions

The project was conceived and supervised by A.J.P. and M.A.B.. T.L.S. performed metagenomic and metatranscriptomic analysis including construction of gene and MAG databases, and statistical analysis including visualization. J.S. supported discussion of MAG metabolism in ecological context. F.B. supported phylogenetic analysis of homologous genes. A.S. and T.L.V.B. supported bioinformatic processing and data visualization. New river sample data including metagenomes and MAGs reported here were provided by M.J.W., J.C.S, A.G., and M.A.B. All authors revised the final version of the manuscript.

## Acknowledgments

We acknowledge the WHONDRS Summer 2019 Sampling (S19S) study and we thank those who participated in the design and implementation of that effort.

## Supplementary Information

**Figure S1:**
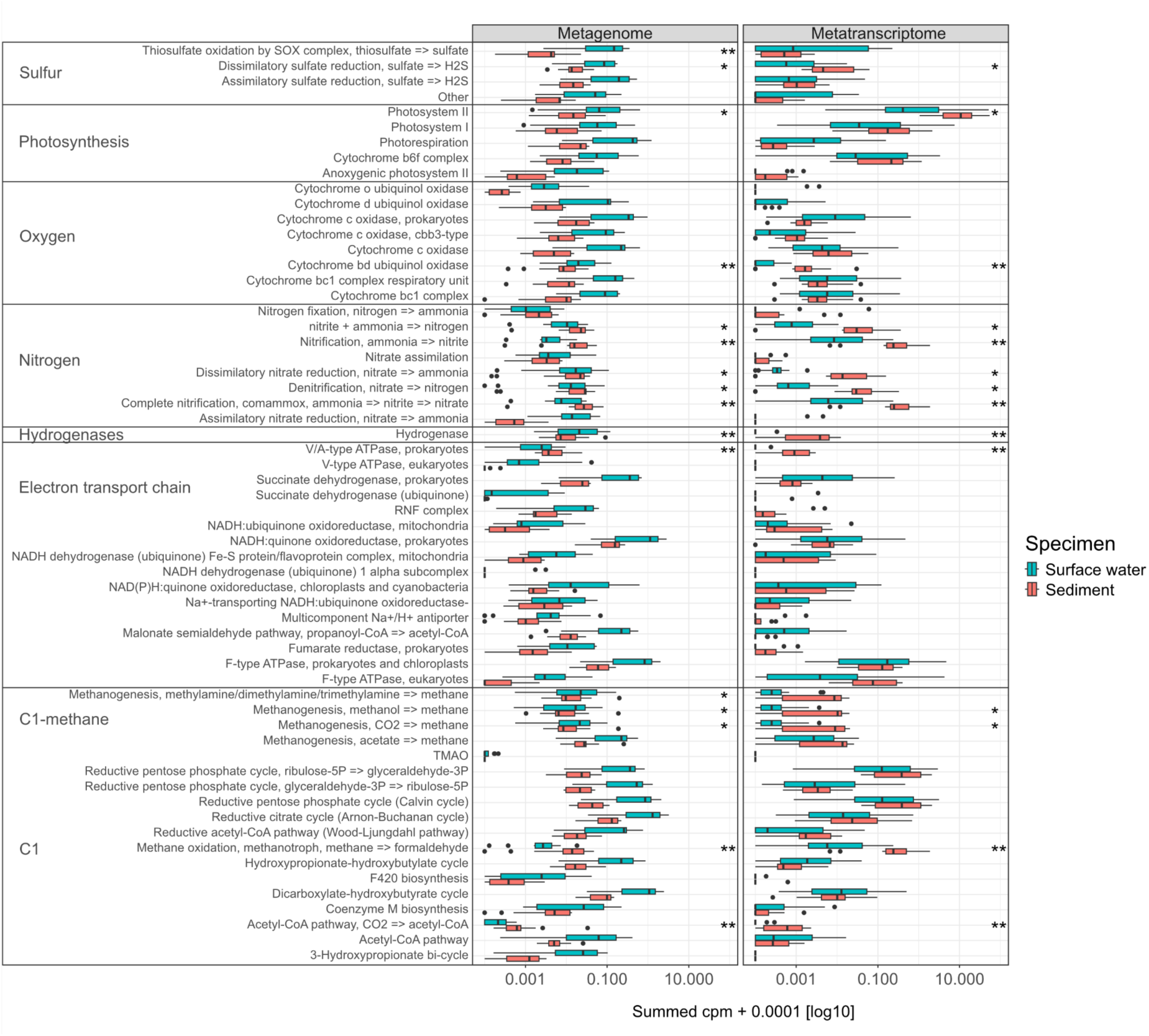
Visualization of the statistical comparison of microbial energy metabolism in sediment and surface water, encoded by the metagenome and expressed by the metatranscriptome. The mean counts per pathway were tested for significant differences between the two specimens (Kruskal-Wallis-Test, n=13); a significantly higher mean is marked by significance indices in the respective column. As support for main Figure 2.

**Figure S2:**
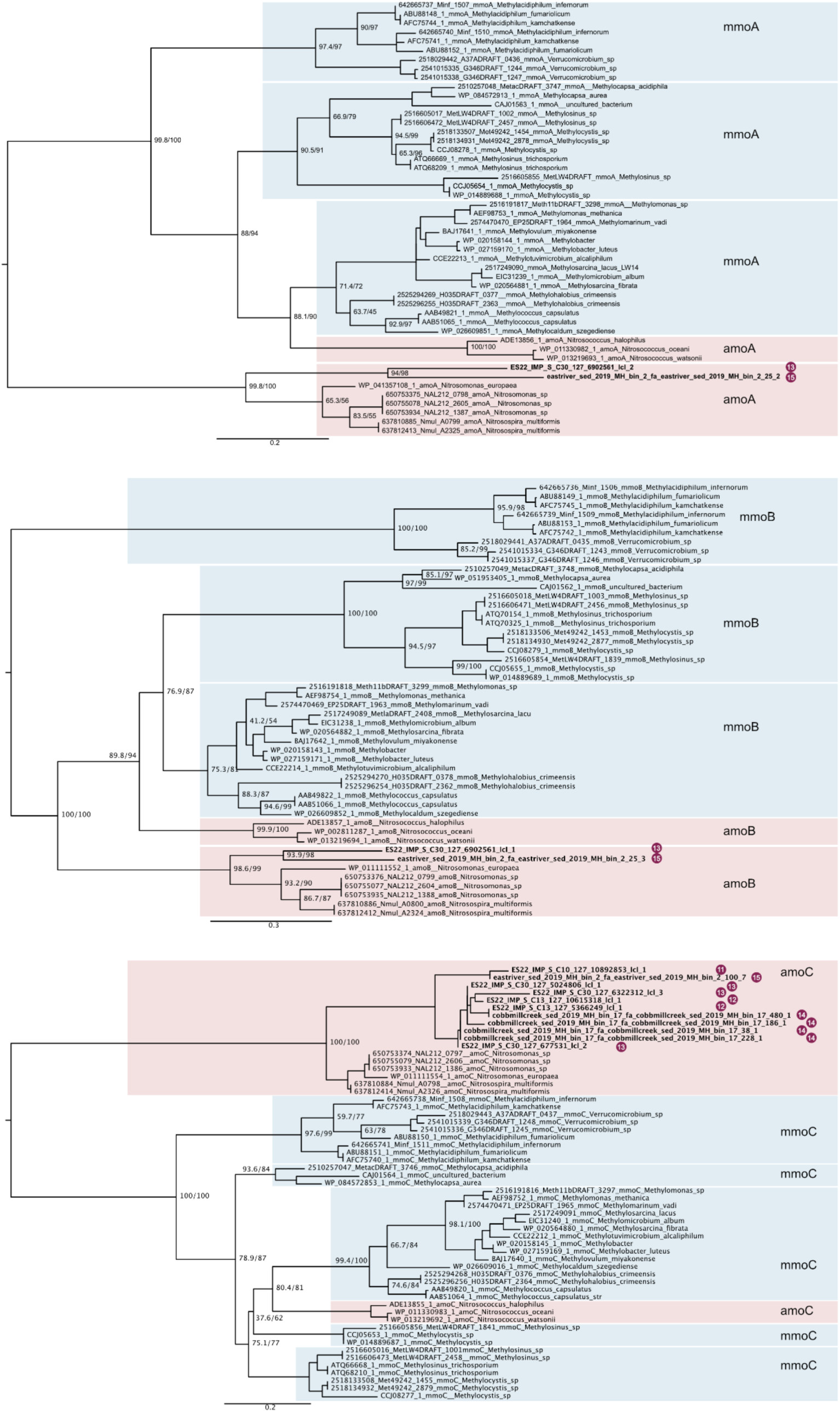
Phylogenetic analysis of homologous genes annotated as methane monooxygenase (mmo) and ammonia monooxygenase (amo), respectively, to decipher the functional annotation of genes retrieved from river metagenomes. The obtained results were used to decide the metabolic capacities of MAGs as depicted in Figure 3. Support corresponds to SH-aLRT support (%) / ultrafast bootstrap support (%).

**Figure S3:**
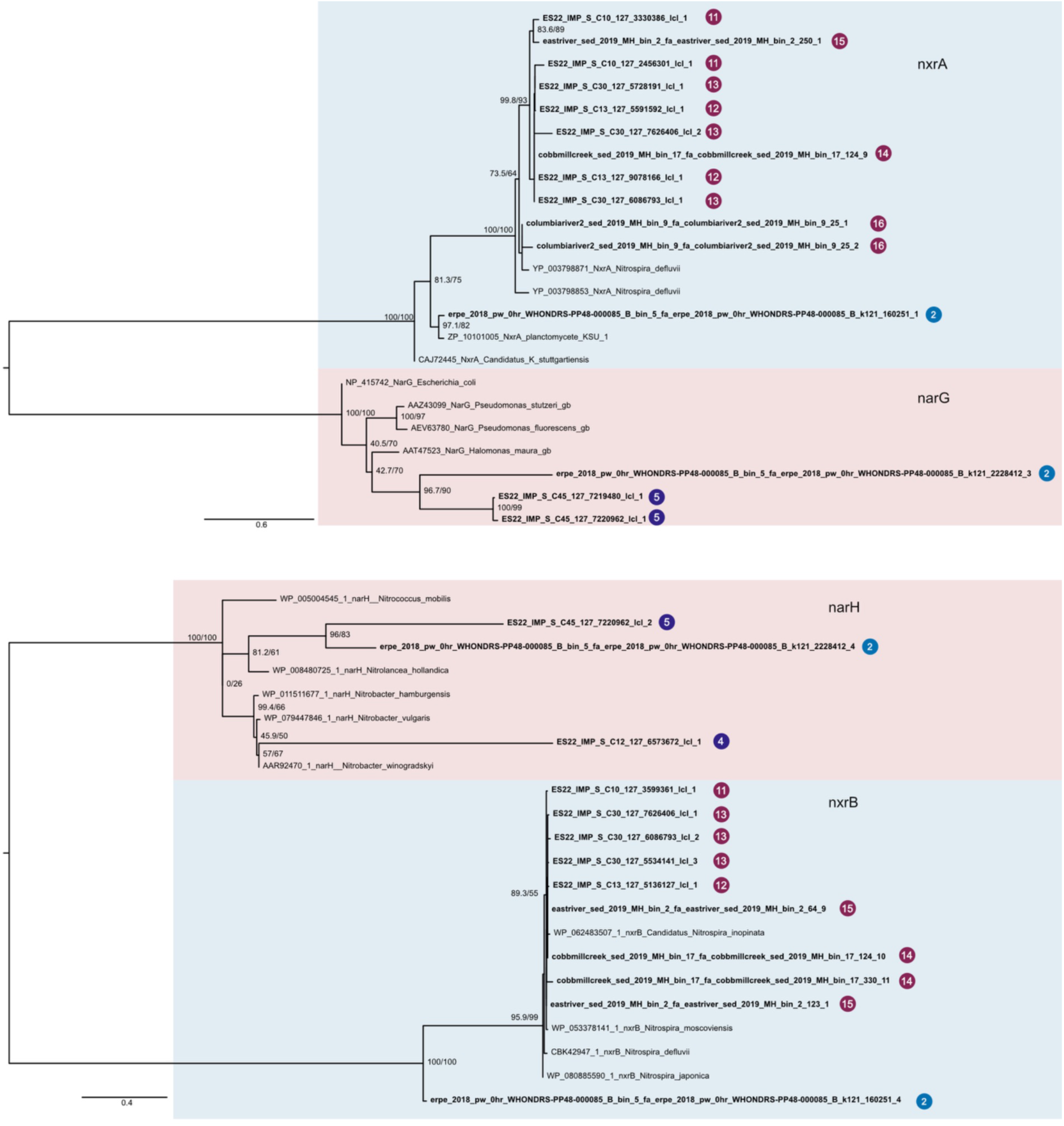
Phylogenetic analysis of homologous genes annotated as nitrate reductase (nar) and nitrite oxidoreductase (nxr), respectively, to decipher proper annotation of homologous genes from river ecosystems. The obtained results were used to decide the metabolic capacities of MAGs as depicted in Figure 3. Support corresponds to SH-aLRT support (%) / ultrafast bootstrap support (%).

**Table S1:** Statistical testing of environmental impact on gene expression of the microbial community structure in stream sediments. Adonis2 was run with 999 permutations and a model including the factors listed below in an additive way based on the marginal effects of the terms as test design.

| Descriptor | Df | SumOfSqs | R<sup>2</sup> | F | Pr(>F) | Sign. |
| --- | --- | --- | --- | --- | --- | --- |
| Temperature | 1 | 0.472 | 0.068 | 1.662 | 0.014 | * |
| Depth | 1 | 0.295 | 0.043 | 1.039 | 0.417 |  |
| Percent total Sand | 1 | 0.439 | 0.064 | 1.545 | 0.039 | * |
| Stream order | 1 | 0.382 | 0.055 | 1.343 | 0.087 | . |
| Nitrogen Percent:<br>Carbon Percent | 1 | 0.339 | 0.049 | 1.194 | 0.221 |  |

**Figure S4 and S5:**
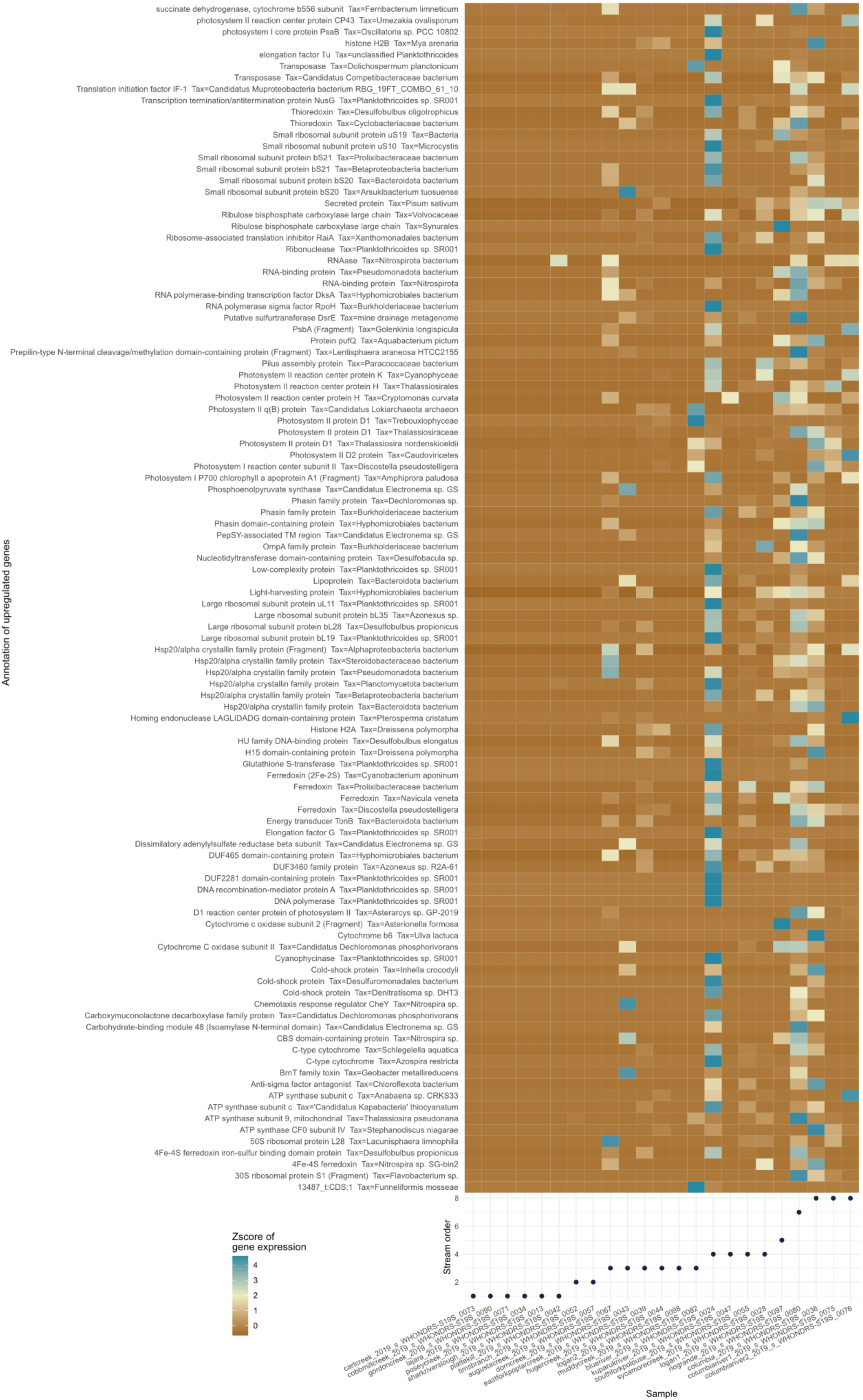
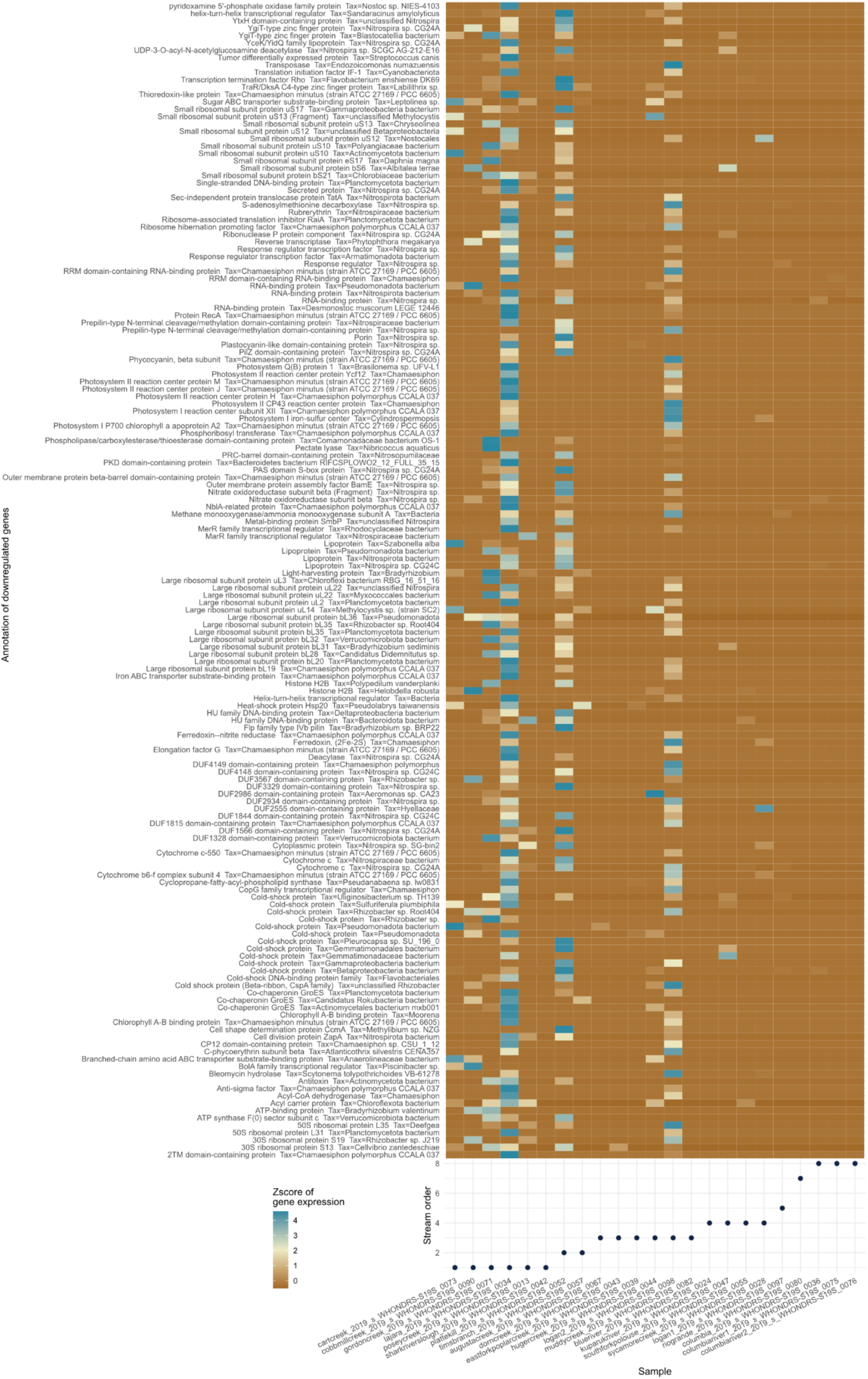
Significantly upregulated (S4) and downregulated (S5) genes with abs(LFC) > 1 and adj. p-value < 0.05 due to stream order increase. Genes annotated as uncharacterized or hypothetical are not shown.

## Notes

### Competing Interest Statement

The authors have declared no competing interest.

## References

1. Henson MW, Hanssen J, Spooner G, Fleming P, Pukonen M, Stahr F, et al. Nutrient dynamics and stream order influence microbial community patterns along a 2914 kilometer transect of the Mississippi River. Limnol Oceanogr. 2018;63:1837–55.

2. Leibowitz SG, Wigington PJ, Schofield KA, Alexander LC, Vanderhoof MK, Golden HE. Connectivity of Streams and Wetlands to Downstream Waters: An Integrated Systems Framework. JAWRA J Am Water Resour Assoc. 2018;54:298–322.

3. Vannote RL, Minshall GW, Cummins KW, Sedell JR, Cushing CE. The River Continuum Concept. Can J Fish Aquat Sci. 1980;37:130–7.

4. Aeschbacher J, Liniger H, Weingartner R. River Water Shortage in a Highland–Lowland System: A Case Study of the Impacts of Water Abstraction in the Mount Kenya Region. Mt Res Dev. 2005;25:155–62.

5. Ferreira V, Albariño R, Larrañaga A, LeRoy CJ, Masese FO, Moretti MS. Ecosystem services provided by small streams: an overview. Hydrobiologia. 2022;850:2501–35.

6. Risse-Buhl U, Anlanger C, Chatzinotas A, Noss C, Lorke A, Weitere M. Near streambed flow shapes microbial guilds within and across trophic levels in fluvial biofilms. Limnol Oceanogr. 2020;65:2261–77.

7. Besemer K, Singer G, Hödl I, Battin TJ. Bacterial Community Composition of Stream Biofilms in Spatially Variable-Flow Environments. Appl Environ Microbiol. 2009;75:7189–95.

8. Jusik S, Szoszkiewicz K, Kupiec JM, Lewin I, Samecka-Cymerman A. Development of comprehensive river typology based on macrophytes in the mountain-lowland gradient of different Central European ecoregions. Hydrobiologia. 2015;745:241–62.

9. Stenger-Kovács C, Tóth L, Tóth F, Hajnal É, Padisák J. Stream order-dependent diversity metrics of epilithic diatom assemblages. Hydrobiologia. 2014;721:67–75.

10. Horton RE. Erosional development of streams and their drainage basins; hydrophysical approach to quantitative morphology. Geol Soc Am Bull. 1945;56:275.

11. Strahler AN. Quantitative analysis of watershed geomorphology. Eos Trans Am Geophys Union. 1957;38:913–20.

12. Alexander RB, Boyer EW, Smith RA, Schwarz GE, Moore RB. The Role of Headwater Streams in Downstream Water Quality1. JAWRA J Am Water Resour Assoc. 2007;43:41–59.

13. Rutherford JC, Marsh NA, Davies PM, Bunn SE. Effects of patchy shade on stream water temperature: how quickly do small streams heat and cool? Mar Freshw Res. 2004;55:737.

14. Hong S, Gim J-S, Kim HG, Cowan PE, Joo G-J. A molecular approach to identifying the relationship between resource use and availability in Eurasian otters (Lutra lutra). Can J Zool. 2019;97:797–804.

15. Whiteside BG, McNatt RM. Fish Species Diversity in Relation to Stream Order and Physicochemical Conditions in the Plum Creek Drainage Basin. Am Midl Nat. 1972;88:90.

16. Fasching C, Akotoye C, Bižić M, Fonvielle J, Ionescu D, Mathavarajah S, et al. Linking stream microbial community functional genes to dissolved organic matter and inorganic nutrients. Limnol Oceanogr. 2020;65:S71–87.

17. Tee HS, Waite D, Lear G, Handley KM. Microbial river-to-sea continuum: gradients in benthic and planktonic diversity, osmoregulation and nutrient cycling. Microbiome. 2021;9:190.

18. Borton MA, McGivern BB, Willi KR, Woodcroft BJ, Mosier AC, Singleton DM, et al. A functional microbiome catalogue crowdsourced from North American rivers. Nature. 2024;637:103–12.

19. Kuglerová L, Hasselquist EM, Sponseller RA, Muotka T, Hallsby G, Laudon H. Multiple stressors in small streams in the forestry context of Fennoscandia: The effects in time and space. Sci Total Environ. 2021;756:143521.

20. Lopes Simedo MB, Pissarra TCT, Mello Martins AL, Lopes MC, Araújo Costa RC, Zanata M, et al. The Assessment of Hydrological Availability and the Payment for Ecosystem Services: A Pilot Study in a Brazilian Headwater Catchment. Water. 2020;12:2726.

21. Meißner T, Sures B, Feld CK. Multiple stressors and the role of hydrology on benthic invertebrates in mountainous streams. Sci Total Environ. 2019;663:841–51.

22. Waite IR, Munn MD, Moran PW, Konrad CP, Nowell LH, Meador MR, et al. Effects of urban multi-stressors on three stream biotic assemblages. Sci Total Environ. 2019;660:1472–85.

23. European Environment Agency. Europe’s state of water 2024: the need for improved water resilience. Luxembourg: Publications Office; 2024.

24. Belletti B, Garcia De Leaniz C, Jones J, Bizzi S, Börger L, Segura G, et al. More than one million barriers fragment Europe’s rivers. Nature. 2020;588:436–41.

25. Grill G, Lehner B, Thieme M, Geenen B, Tickner D, Antonelli F, et al. Mapping the world’s free-flowing rivers. Nature. 2019;569:215–21.

26. Griffith MB, Norton SB, Alexander LC, Pollard AI, LeDuc SD. The effects of mountaintop mines and valley fills on the physicochemical quality of stream ecosystems in the central Appalachians: A review. Sci Total Environ. 2012;417–418:1–12.

27. Růžičková S, Schenková J, Weissová V, Helešic J. Environmental impact of heated mining waters on clitellate (Annelida: Clitellata) assemblages. Biologia (Bratisl). 2014;69:1179–89.

28. Fodelianakis S, Washburne AD, Bourquin M, Pramateftaki P, Kohler TJ, Styllas M, et al. Microdiversity characterizes prevalent phylogenetic clades in the glacier-fed stream microbiome. ISME J. 2022;16:666–75.

29. Kohler TJ, Bourquin M, Peter H, Yvon-Durocher G, Sinsabaugh RL, Deluigi N, et al. Global emergent responses of stream microbial metabolism to glacier shrinkage. Nat Geosci. 2024;17:309–15.

30. David GM, Pimentel IM, Rehsen PM, Vermiert A-M, Leese F, Gessner MO. Multiple stressors affecting microbial decomposer and litter decomposition in restored urban streams: Assessing effects of salinization, increased temperature, and reduced flow velocity in a field mesocosm experiment. Sci Total Environ. 2024;943:173669.

31. Romero F, Acuña V, Sabater S. Multiple Stressors Determine Community Structure and Estimated Function of River Biofilm Bacteria. Schaffner DW, editor. Appl Environ Microbiol. 2020;86:e00291–20.

32. U.S. Environmental Protection Agency. National Rivers and Streams Assessment: The Third Collaborative Survey. U.S. Environmental Protection Agency, Office of Water and Office of Research and Development; 2024. Report No.: EPA 841-R-22-004.

33. Lewandowski J, Arnon S, Banks E, Batelaan O, Betterle A, Broecker T, et al. Is the Hyporheic Zone Relevant beyond the Scientific Community? Water. 2019;11:2230.

34. Nelson SM, Roline RA. Effects of multiple stressors on hyporheic invertebrates in a lotic system. Ecol Indic. 2003;3:65–79.

35. Hancock PJ. Human Impacts on the Stream-Groundwater Exchange Zone. Environ Manage. 2002;29:763–81.

36. Doloiras-Laraño AD, Serrana JM, Takahashi S, Takemon Y, Watanabe K. Short-term influences of flow alteration on microbial community structure and putative metabolic functions in gravel bar hyporheic zones. Front Environ Sci. 2023;11:1205561.

37. Fischer H, Kloep F, Wilzcek S, Pusch MT. A River’s Liver – Microbial Processes within the Hyporheic Zone of a Large Lowland River. Biogeochemistry. 2005;76:349–71.

38. Wang Y, Wang Y, Shang J, Wang L, Li Y, Wang Z, et al. Redox gradients drive microbial community assembly patterns and molecular ecological networks in the hyporheic zone of effluent-dominated rivers. Water Res. 2024;248:120900.

39. Gerbersdorf SU, Jancke T, Westrich B, Paterson DM. Microbial stabilization of riverine sediments by extracellular polymeric substances. Geobiology. 2007;6:57–69.

40. Ezzat L, Fodelianakis S, Kohler TJ, Bourquin M, Brandani J, Busi SB, et al. Benthic Biofilms in Glacier-Fed Streams from Scandinavia to the Himalayas Host Distinct Bacterial Communities Compared with the Streamwater. Rudi K, editor. Appl Environ Microbiol. 2022;88:e00421–22.

41. Rodríguez-Ramos JA, Oliverio A, Borton MA, Danczak R, Mueller BM, Schulz H, et al. Spatial and temporal metagenomics of river compartments reveals viral community dynamics in an urban impacted stream. Front Microbiomes. 2023;2:1199766.

42. Piggott JJ, Salis RK, Lear G, Townsend CR, Matthaei CD. Climate warming and agricultural stressors interact to determine stream periphyton community composition. Glob Change Biol. 2015;21:206–22.

43. Beermann AJ, Elbrecht V, Karnatz S, Ma L, Matthaei CD, Piggott JJ, et al. Multiple-stressor effects on stream macroinvertebrate communities: A mesocosm experiment manipulating salinity, fine sediment and flow velocity. Sci Total Env. 2018;610–611:961–71.

44. Graupner N, Röhl O, Jensen M, Beisser D, Begerow D, Boenigk J. Effects of short-term flooding on aquatic and terrestrial microeukaryotic communities: a mesocosm approach. Aquat Microb Ecol. 2017;80:257–72.

45. Burdon FJ, Bai Y, Reyes M, Tamminen M, Staudacher P, Mangold S, et al. Stream microbial communities and ecosystem functioning show complex responses to multiple stressors in wastewater. Glob Change Biol. 2020;26:6363–82.

46. Subirats J, Triadó-Margarit X, Mandaric L, Acuña V, Balcázar JL, Sabater S, et al. Wastewater pollution differently affects the antibiotic resistance gene pool and biofilm bacterial communities across streambed compartments. Mol Ecol. 2017;26:5567–81.

47. Stegen JC, Goldman AE. WHONDRS: a Community Resource for Studying Dynamic River Corridors. mSystems [Internet]. 2018 [cited 2024 Nov 20];3. Available from: https://journals.asm.org/doi/10.1128/msystems.00151-18

48. Thoetkiattikul H, Mhuantong W, Pinyakong O, Wisawapipat W, Yamazoe A, Fujita N, et al. Culture-independent study of bacterial communities in tropical river sediment. Biosci Biotechnol Biochem. 2017;81:200–9.

49. Anantharaman K, Brown CT, Hug LA, Sharon I, Castelle CJ, Probst AJ, et al. Thousands of microbial genomes shed light on interconnected biogeochemical processes in an aquifer system. Nat Commun. 2016;7:13219.

50. Rodríguez-Ramos JA, Borton MA, McGivern BB, Smith GJ, Solden LM, Shaffer M, et al. Genome-Resolved Metaproteomics Decodes the Microbial and Viral Contributions to Coupled Carbon and Nitrogen Cycling in River Sediments. Hug LA, editor. mSystems. 2022;7:e00516–22.

51. Brasseur MV, Buchner D, Mack L, Schreiner VC, Schäfer RB, Leese F, et al. Multiple stressor effects of insecticide exposure and increased fine sediment deposition on the gene expression profiles of two freshwater invertebrate species. Environ Sci Eur. 2023;35:81.

52. Elbrecht V, Beermann AJ, Goessler G, Neumann J, Tollrian R, Wagner R, et al. Multiple-stressor effects on stream invertebrates: a mesocosm experiment manipulating nutrients, fine sediment and flow velocity. Freshw Biol. 2016;61:362–75.

53. Madge Pimentel I, Rehsen PM, Beermann AJ, Leese F, Piggott JJ, Schmuck S. An automated modular heating solution for experimental flow-through stream mesocosm systems. Limnol Oceanogr Methods. 2023;lom3.10596.

54. Nuy JK, Lange A, Beermann AJ, Jensen M, Elbrecht V, Röhl O, et al. Responses of stream microbes to multiple anthropogenic stressors in a mesocosm study. Sci Total Environ. 2018;633:1287–301.

55. Stach TL, Deep A, Madge Pimentel I, Buchner D, Borton MA, Soares A, et al. Complex compositional and cellular response of river sediment microbiomes to multiple anthropogenic stressors. bioRxiv. 2024;2024.06.07.597903.

56. Albertsen M, Hugenholtz P, Skarshewski A, Nielsen KL, Tyson GW, Nielsen PH. Genome sequences of rare, uncultured bacteria obtained by differential coverage binning of multiple metagenomes. Nat Biotechnol. 2013;31:533–8.

57. Haryono MAS, Law YY, Arumugam K, Liew LC-W, Nguyen TQN, Drautz-Moses DI, et al. Recovery of High Quality Metagenome-Assembled Genomes From Full-Scale Activated Sludge Microbial Communities in a Tropical Climate Using Longitudinal Metagenome Sampling. Front Microbiol. 2022;13:869135.

58. Madge Pimentel I, Baikova D, Buchner D, Burfeid Castellanos A, David GM, Deep A, et al. Assessing the response of an urban stream ecosystem to salinization under different flow regimes. Sci Total Environ. 2024;926:171849.

59. Goldman AE, Arnon S, Bar-Zeev E, Chu RK, Danczak RE, Daly RA, et al. WHONDRS Summer 2019 Sampling Campaign: Global River Corridor Sediment FTICR-MS, Dissolved Organic Carbon, Aerobic Respiration, Elemental Composition, Grain Size, Total Nitrogen and Organic Carbon Content, Bacterial Abundance, and Stable Isotopes (v8) [Internet]. Environmental System Science Data Infrastructure for a Virtual Ecosystem; River Corridor and Watershed Biogeochemistry SFA; 2020 [cited 2024 Nov 20]. Available from: https://www.osti.gov/servlets/purl/1729719/

60. Joshi N, Fass J. Sickle: A sliding-window, adaptive, quality-based trimming tool for FastQ files [Internet]. 2011. Available from: https://github.com/najoshi/sickle

61. Peng Y, Leung HCM, Yiu SM, Chin FYL. IDBA-UD: a de novo assembler for single-cell and metagenomic sequencing data with highly uneven depth. Bioinformatics. 2012;28:1420–8.

62. Li D, Luo R, Liu C-M, Leung C-M, Ting H-F, Sadakane K, et al. MEGAHIT v1.0: A fast and scalable metagenome assembler driven by advanced methodologies and community practices. Methods. 2016;102:3–11.

63. Kang DD, Li F, Kirton E, Thomas A, Egan R, An H, et al. MetaBAT 2: an adaptive binning algorithm for robust and efficient genome reconstruction from metagenome assemblies. PeerJ. 2019;7:e7359.

64. Olm MR, Brown CT, Brooks B, Banfield JF. dRep: a tool for fast and accurate genomic comparisons that enables improved genome recovery from metagenomes through de-replication. ISME J. 2017;11:2864–8.

65. Chklovski A, Parks DH, Woodcroft BJ, Tyson GW. CheckM2: a rapid, scalable and accurate tool for assessing microbial genome quality using machine learning. Nat Methods. 2023;20:1203–12.

66. Chaumeil P-A, Mussig AJ, Hugenholtz P, Parks DH. GTDB-Tk: a toolkit to classify genomes with the Genome Taxonomy Database. Bioinformatics. 2020;36:1925–7.

67. Hyatt D, Chen G-L, LoCascio PF, Land ML, Larimer FW, Hauser LJ. Prodigal: prokaryotic gene recognition and translation initiation site identification. BMC Bioinformatics. 2010;11:119.

68. Langmead B, Salzberg SL. Fast gapped-read alignment with Bowtie 2. Nat Methods. 2012;9:357–9.

69. Shaffer M, Borton MA, McGivern BB, Zayed AA, La Rosa SL, Solden LM, et al. DRAM for distilling microbial metabolism to automate the curation of microbiome function. Nucleic Acids Res. 2020;48:8883–900.

70. Woodcroft BJ, Aroney STN, Zhao R, Cunningham M, Mitchell JAM, Blackall L, et al. SingleM and Sandpiper: Robust microbial taxonomic profiles from metagenomic data. bioRxiv. 2024;2024.01.30.578060.

71. Putri GH, Anders S, Pyl PT, Pimanda JE, Zanini F. Analysing high-throughput sequencing data in Python with HTSeq 2.0. Boeva V, editor. Bioinformatics. 2022;38:2943–5.

72. Yunshun Chen AL. edgeR [Internet]. Bioconductor; 2017. Available from: https://bioconductor.org/packages/edgeR

73. Xu S, Dai Z, Guo P, Fu X, Liu S, Zhou L, et al. ggtreeExtra: Compact Visualization of Richly Annotated Phylogenetic Data. Mol Biol Evol. 2021;38:4039–42.

74. Campitelli E. ggnewscale: Multiple Fill and Colour Scales in “ggplot2” [Internet]. 2024. Available from: https://CRAN.R-project.org/package=ggnewscale

75. Katoh K, Standley DM. MAFFT Multiple Sequence Alignment Software Version 7: Improvements in Performance and Usability. Mol Biol Evol. 2013;30:772–80.

76. Capella-Gutiérrez S, Silla-Martínez JM, Gabaldón T. trimAl: a tool for automated alignment trimming in large-scale phylogenetic analyses. Bioinformatics. 2009;25:1972–3.

77. Minh BQ, Schmidt HA, Chernomor O, Schrempf D, Woodhams MD, Von Haeseler A, et al. IQ-TREE 2: New Models and Efficient Methods for Phylogenetic Inference in the Genomic Era. Teeling E, editor. Mol Biol Evol. 2020;37:1530–4.

78. Rochman FF, Kwon M, Khadka R, Tamas I, Lopez-Jauregui AA, Sheremet A, et al. Novel copper-containing membrane monooxygenases (CuMMOs) encoded by alkane-utilizing Betaproteobacteria. ISME J. 2020;14:714–26.

79. Steinegger M, Söding J. MMseqs2 enables sensitive protein sequence searching for the analysis of massive data sets. Nat Biotechnol. 2017;35:1026–8.

80. Toyoda JG, Goldman AE, Arnon S, Bar-Zeev E, Chu RK, Danczak RE, et al. WHONDRS Summer 2019 Sampling Campaign: Global River Corridor Surface Water FTICR-MS, NPOC, TN, Anions, Stable Isotopes, Bacterial Abundance, and Dissolved Inorganic Carbon (v6) [Internet]. Environmental System Science Data Infrastructure for a Virtual Ecosystem; River Corridor and Watershed Biogeochemistry SFA; 2020 [cited 2025 Jan 22]. Available from: https://www.osti.gov/servlets/purl/1603775/

81. Chen J, Zhang X, Yang L, Zhang L. GUniFrac: Generalized UniFrac Distances, Distance-Based Multivariate Methods and Feature-Based Univariate Methods for Microbiome Data Analysis [Internet]. 2023. Available from: https://CRAN.R-project.org/package=GUniFrac

82. Oksanen J, Blanchet FG, Kindt R, Legendre P, Minchin P, O’Hara R, et al. Vegan: Community Ecology Package. R Package Version. 2.0-10. CRAN. 2013;

83. Quensen J, Simpson G, Oksanen J. ggordiplots: Make “ggplot2” Versions of Vegan’s Ordiplots [Internet]. 2024. Available from: https://CRAN.R-project.org/package=ggordiplots

84. Simpson GL. permute: Functions for Generating Restricted Permutations of Data [Internet]. 2022. Available from: https://CRAN.R-project.org/package=permute

85. Gehlenborg N. UpSetR: A More Scalable Alternative to Venn and Euler Diagrams for Visualizing Intersecting Sets [Internet]. 2019. Available from: https://CRAN.R-project.org/package=UpSetR

86. Buchfink B, Reuter K, Drost H-G. Sensitive protein alignments at tree-of-life scale using DIAMOND. Nat Methods. 2021;18:366–8.

87. Bornemann TLV, Adam PS, Turzynski V, Schreiber U, Figueroa-Gonzalez PA, Rahlff J, et al. Genetic diversity in terrestrial subsurface ecosystems impacted by geological degassing. Nat Commun. 2022;13:284.

88. Suzek BE, Huang H, McGarvey P, Mazumder R, Wu CH. UniRef: comprehensive and non-redundant UniProt reference clusters. Bioinformatics. 2007;23:1282–8.

89. Wickham H. ggplot2: elegant graphics for data analysis. Second edition. Switzerland: Springer; 2016.

90. Gerson JR, Hinckley ES. It Is Time to Develop Sustainable Management of Agricultural Sulfur. Earths Future. 2023;11:e2023EF003723.

91. Turner RE, Rabalais NN. Linking Landscape and Water Quality in the Mississippi River Basin for 200 Years. BioScience. 2003;53:563.

92. Xia X, Zhang S, Li S, Zhang L, Wang G, Zhang L, et al. The cycle of nitrogen in river systems: sources, transformation, and flux. Environ Sci Process Impacts. 2018;20:863–91.

93. Zhang D, Liu F, Al MA, Yang Y, Yu H, Li M, et al. Nitrogen and sulfur cycling and their coupling mechanisms in eutrophic lake sediment microbiomes. Sci Total Environ. 2024;928:172518.

94. Van Kessel MAHJ, Speth DR, Albertsen M, Nielsen PH, Op Den Camp HJM, Kartal B, et al. Complete nitrification by a single microorganism. Nature. 2015;528:555–9.

95. Daims H, Lebedeva EV, Pjevac P, Han P, Herbold C, Albertsen M, et al. Complete nitrification by Nitrospira bacteria. Nature. 2015;528:504–9.

96. Bayer B, Saito MA, McIlvin MR, Lücker S, Moran DM, Lankiewicz TS, et al. Metabolic versatility of the nitrite-oxidizing bacterium Nitrospira marina and its proteomic response to oxygen-limited conditions. ISME J. 2021;15:1025–39.

97. Lücker S, Wagner M, Maixner F, Pelletier E, Koch H, Vacherie B, et al. A Nitrospira metagenome illuminates the physiology and evolution of globally important nitrite-oxidizing bacteria. Proc Natl Acad Sci. 2010;107:13479–84.

98. Wang Y, Bott C, Nerenberg R. Sulfur-based denitrification: Effect of biofilm development on denitrification fluxes. Water Res. 2016;100:184–93.

99. Wang Z, He S, Huang J, Zhou W, Chen W. Comparison of heterotrophic and autotrophic denitrification processes for nitrate removal from phosphorus-limited surface water. Environ Pollut. 2018;238:562–72.

100. Lücker S, Nowka B, Rattei T, Spieck E, Daims H. The Genome of Nitrospina gracilis Illuminates the Metabolism and Evolution of the Major Marine Nitrite Oxidizer. Front Microbiol. 2013;4.

101. Cheng B, Xia R, Zhang Y, Yang Z, Hu S, Guo F, et al. Characterization and causes analysis for algae blooms in large river system. Sustain Cities Soc. 2019;51:101707.

102. Kaushal SS, Likens GE, Jaworski NA, Pace ML, Sides AM, Seekell D, et al. Rising stream and river temperatures in the United States. Front Ecol Environ. 2010;8:461–6.

103. Vos M, Hering D, Gessner MO, Leese F, Schäfer RB, Tollrian R, et al. The Asymmetric Response Concept explains ecological consequences of multiple stressor exposure and release. Sci Total Environ. 2023;872:162196.

104. Xu M, Wang Z, Pan B, Zhao N. Distribution and species composition of macroinvertebrates in the hyporheic zone of bed sediment. Int J Sediment Res. 2012;27:129–40.

105. Bianchin MS, Smith L, Beckie RD. Defining the hyporheic zone in a large tidally influenced river. J Hydrol. 2011;406:16–29.

106. Peters K, Grantz SF, Kiesel J, Lewandowski J, Fohrer N. Hyporheic exchange flows in a mountainous river catchment identified by distributed temperature sensing. River Res Appl. 2024;40:1417–31.

107. Hug LA, Hatzenpichler R, Moraru C, Soares A, Meyer F, Heyder A, et al. A roadmap for fair reuse of public microbiome data. bioRxiv. 2024;2024.06.21.599698.

108. Wilkinson MD, Dumontier M, Aalbersberg IjJ, Appleton G, Axton M, Baak A, et al. The FAIR Guiding Principles for scientific data management and stewardship. Sci Data. 2016;3:160018.

